# Mouse model of the human serotonin transporter-linked polymorphic region

**DOI:** 10.1101/556092

**Authors:** Lukasz Piszczek, Simone Memoli, Angelo Raggioli, José Viosca, Jeanette Rientjes, Philip Hublitz, Weronika Czaban, Anna Wyrzykowska, Cornelius Gross

**Affiliations:** Epigenetics and Neurobiology Unit, European Molecular Biology Laboratory, EMBL Rome, Monterotondo, Italy; Research Institute of Molecular Pathology, Vienna, Austria; Promotion of Health and Biomedical Research in the Valencian Region (FISABIO), Valencia, Spain; Monash Genome Modification Platform (MGMP), Monash University, Clayton, Australia; MRC Weatherall Institute of Molecular Medicine, University of Oxford, UK

**Keywords:** (4-6) 5-HTT, 5-HTT-LPR, serotonin, mouse models

## Abstract

Genetic factors play a significant role in risk for mood and anxiety disorders. Polymorphisms in genes that regulate the brain monoamine systems, such as catabolic enzymes and transporters, are attractive candidates for being risk factors for emotional disorders given the weight of evidence implicating monoamines involvement in these conditions. Several common genetic variants have been identified in the human serotonin transporter (5-HTT) gene, including a repetitive sequence located in the promoter region of the locus called the serotonin transporter-linked polymorphic region (5-HTT-LPR). This polymorphism has been associated with a number of mental traits in both humans and primates, including depression, neuroticism, and harm avoidance. Some, but not all studies found a link between the polymorphism and 5-HTT levels, leaving open the question of whether the polymorphism affects risk for mental traits via changes in 5-HTT expression. To investigate the impact of the polymorphism on gene expression, serotonin homeostasis, and behavioural traits we set out to develop a mouse model of the human 5-HTT- LPR. Here we describe the creation and characterization of a set of mouse lines with single copy human transgenes carrying the short and long 5-HTT-LPR variants.

## Introduction

Disturbances in the serotonin (5-hydroxytryptamine, 5-HT) system have been implicated in the pathophysiology of a wide variety of mental disorders, including anxiety, depression, aggression, addiction, and suicide. The serotonin transporter (5-HTT, *SLC6A4*) regulates the spatio-temporal fine-tuning of 5-HT signaling by mediating the re-uptake of serotonin from the synaptic cleft. Drugs that block 5-HTT prolong the synaptic lifetime of serotonin and boost the activation of its receptor targets are effective antidepressants. At the same time, genetic variants in 5-HTT that reduce its expression or functionality are associated with increased risk for mood disorders and its associated traits (Malison *et al*, 1998, Little *et al*, 1998, Purselle & Nemeroff, 2003). The apparent opposite effects of pharmacological and genetic loss-of-function manipulations of 5-HTT remain unexplained, but have been proposed to depend either on the engagement of negative feedback mechanisms on serotonin neuron firing triggered by acute pharmacological inhibition or on the impact of genetic variations on brain developmental trajectories (Ansorge et al, 2004, Homberg et al, 2010, Oh et al, 2009, Popa et al, 2008).

Several common genetic variants have been identified in the human 5-HTT gene. Of these, the most extensively studied is a polymorphic repetitive sequence located upstream of the transcription start site called the serotonin transporter-linked polymorphic region (5-HTT-LPR; Lesch et al., 1997; Murphy *et al*, 2004). This promoter region is composed of repetitive nucleotide sequences each 20 to 23 base pairs long, with the most frequent alleles called the “short” allele (S), containing 14 repeats, and the “long” allele (L), containing 16 repeats (Lesch et al., 1997; Colucci et al, 2008). The most common sub-variant of the L allele is called 16A, while that of the S allele is called 14A (Nakamura et al, 2000). Interestingly, the frequency of the 5-HTT S and L variants differs significantly across human populations (Noskova et al, 2008). Moreover, the 5-HTT-LPR is only present in humans and higher primates, but not in prosimians and rodents (Lesch et al, 1997). The lack of a suitable mouse model of 5-HTT-LPR has hampered the understanding of its molecular, cellular, and physiological impact on mental traits.

Several studies found a link between the 5-HTT-LPR S allele, the presence of environmental adversity, and increased risk for major depression. Individuals homozygous for the S allele of the polymorphism were more susceptible to the negative effects of life stressors in early childhood than those homozygous for the L allele, resulting in an increased risk for major depression (Caspi et al, 2003, Kaufman et al, 2004, Kendler et al, 2005, but see Munafò et al, 2008 and Risch et al, 2009). In the absence of identifiable stress events no increased risk for depression was observed, suggesting that the major mechanisms by which 5-HTT-LPR moderates mental traits may be via its effect on the impact of environmental stressors. Nevertheless, some studies that did not quantify environmental stressors did find increased anxiety-related personality traits in S allele carriers, including increased neuroticism and harm avoidance (Lesch et al, 1996; Greenberg et al, 2000). Moreover, several studies have shown differences in S allele carriers in steady-state and event-related neural activity in cortex and amygdala as measured by functional magnetic resonance imaging (fMRI), a finding that was more pronounced in depressed subjects (Hariri & Holmes, 2006; Canli et al, 2005; Heinz et al, 2005; Dannlowski et al, 2007; Lau et al, 2009; Pezawas et al, 2005).

Although some data demonstrates that the 5-HTT-LPR S allele is associated with reduced expression of the 5-HTT gene in human lymphocytes (Lesch et al, 1996; Heils et al, 1996, Hranilovic et al, 2004; Lima et al, 2005), the magnitude, direction, and time course of this effect in the human brain remains unclear. Positron emission tomography (PET) imaging studies in the living brain have shown a correlation between the S allele and decreased 5-HTT ligand binding levels (Reimold et al, 2007), however, others reported no such correlation (Parsey et al, 2006; Shioe et al, 2003). Together, these studies suggest that the effect of 5-HTT-LPR on gene expression may be cell-type and/or time-dependent, with greater differences possibly occurring at early developmental stages. Such an effect would be consistent with animal studies in which pharmacological blockade of 5-HTT during the first postnatal weeks is sufficient to alter anxiety-related behavior measured later in adulthood (Ansorge et al, 2004).

To facilitate the study of the molecular, cellular, and physiological consequences of 5-HTT-LPR variants we developed a set of transgenic mouse lines carrying the human 5-HTT locus with either the 5-HTT-LPR-16A (L, long) or 14A (S, short) alleles. Variants were introduced into the human 5-HTT locus into a bactrerial artificial chromosome (BAC) using recombineering in bacteria. The inclusion of both, fluorescence and luciferase reporter genes into the 5-HTT gene open reading frame facilitated the assessment of transgene spatial expression and quantitative comparison at low expression levels, respectively. To allow for direct comparisons of S and L allele function in mouse, we inserted the human 5-HTT locus at the same position in each line using recombination-mediated cassette exchange (RMCE). Here we present data describing the expression of the S and L variant human transgenes in mouse.

## Materials and methods

### Targeting constructs

The FRT-PGK-em7-Keo-FRT cassette was amplified using high-fidelity PCR from pPKG-em7-Keo-FRT and used for TOPO-TA cloning into the pCRII plasmid (TOPO TA Cloning Kit Dual Promoter, Invitrogen). SYFP2 coding sequence (from pSYFP2-C1, Kremers et al, 2006) was amplified via high-fidelity PCR and inserted in-frame along with a firefly luciferase coding sequence (amplified from psiCHECK-2, Promega) with a P2A sequence (Holst et al, 2006) upstream of the firefly luciferase gene. The ends of a human BAC (RP11-126E3, Chori BACPAC Resources) containing a genomic fragment harboring the entire coding region of the 5-HTT gene (*SLC6A4*) was trimmed using bacterial recombineering to produce a 62 kb fragment starting at the end of the flanking BLMH gene and ending at the stop of the CCDC55 gene. The resulting P2A-hLuc-T2A-SYFP-FRT-PGK-Em7-Keo-FRT construct was then inserted to replace the stop codon of the trimmed human BAC (clone 11A6) by recombineering, resulting in RP11-126E3/P2A-hLuc-T2A-SYFP-FKF, called RP11-16A for short.

Several attempts at using bacterial recombineering with negative selection (SacB or Rpsl) to convert RP11-16A into RP11-14A without altering the surrounding sequence failed because positive clones invariably carried large deletions of the BAC and we decided to use recombineering with positive selection followed by removal of the selectable marker via flanking rox recombinase sites. First, a 16A-Rox BAC was constructed by recombining a rox-Zeo-rox cassette into RP11-16A 116 bp upstream of 5-HTT-LPR, after which the Zeo cassette was removed by Dre-mediated excision. Next, a 14A-Rox construct was created by recombineering the rox-Zeo-rox cassette into the previously created pBSK-14A plasmid containing the 14A sequence from the human RP-11-104B7 BAC (Chori BACPAC Resources). Finally, the resulting product was used for recombining it into the previously created RP11-AMP BAC to exchange the AMP-SacB cassette and produce RP11-14A-rox-Zeo-rox. Ultimately, the Zeo cassette was removed by Dre-mediated excision to create RP11-14A-rox.

### Cell line targeting

The RMCE acceptor ES cell line (CGL1.10E) was kindly provided by Haydn Prosser (Sanger Institute, Hinxton, UK). ES cells were cultivated on mitomycin C treated SNL Feeder Cells (kindly provided by Haydn Prosser). The targeting protocol was adapted from a previous published method for RMCE (Prosser *et al*., 2008). Briefly, 10.5 million trypsinized cells were mixed with 10 μg of CsCl purified BAC and 25 μg of pCAGGS-Cre (kindly provided by Olga Ermakova, CNR) in a 0.4 μm cuvette (Bio-Rad) and electroporated (230 V, 500 μF). Cells were collected, centrifuged (5 min, 1000 rpm) and plated on a 10 cm dish. Four to five such electroporations were performed per construct to ensure the recovery of sufficient number of positive clones. After 24 hrs, 200 μg/ml G418 (Invitrogen) was added to the medium and the selection continued for 2-3 days, after which 10 μM 6-TG (6-thioguanine or 2- amino-6-mercaptopurine, Sigma-Aldrich) with 200 μg/ml G418 selection was carried out for an additional 6-8 days. Resistant clones were picked in a SNL-seeded 96-well plate and frozen down after reaching confluence. During the picking of clones a replica plate with 0.1% gelatin-coated wells was prepared for Southern Blot screening. For blastocyst injection positive clones were thawed onto a 48-well plate, passaged on 12-well plates, and expanded on 6-well plates before transfer on ice to the EMBL Transgenic Facility.

### Mouse lines

The generation and characterization of the *Tph2*::rLuc BAC transgenic line was published elsewhere (Mlinar et al. 2016). This line was induced and maintained on an inbred FVB/J genetic background, crossed to the created 5-HTT-LPR lines for testing on an F1 hybrid genetic background.

### Animal husbandry

Animals were housed in groups of two to four per cage with free access to food and water. Animals were maintained on a 12:12 light/dark schedule (lights on at 7:00, off at 19:00). All mice were handled according to protocols approved by the Italian Ministry of Health (#137/2011-B, #231/2011-B, #541/2015-PR) and commensurate with NIH guidelines for the ethical treatment of animal.

### Southern blot

Mouse embryonic stem cells (ES) were grown to confluency, washed twice with 1xPBS and lysed in buffer (10 mM Tris pH 7.5, 10 mM EDTA, 10 mM NaCl, 0.5% sarcosyl, 1 mg/ml Proteinase K). The plate was incubated at 56° C overnight in a humid atmosphere. DNA was ethanol-precipitated and resuspended in digestion buffer (1x restriction buffer, 1 mM spermidine, 1 mM DTT, 100 μg/ml BSA, 50 μg/ml RNse A, 50-80 U EcoRI enzyme). The plate was incubated overnight at 37° C in a humid atmosphere. The following day, DNA digest products were resolved by electrophoresis and DNA was transferred according to standard protocol (Southern, 2006) in alkaline solution (0.4 M NaOH, 1.5 M NaCl) onto a nylon membrane (Hybond-N+, Amersham). After transfer, the membrane was washed 3−15 min in 2xSSC, air-dried for 1 hour and cross-linked (150 mJ, UV-Crosslinker, Vilber Lourmat). Southern blot was performed in hybridization buffer (0.5 M Phosphate Buffer, 1mM EDTA, 3% BSA, 5% SDS) with 25 ng of radioactively labeled probe ([α-32P]-dGTP, PerkinElmer) according to manufacturer instructions (Random Primers DNA Labeling System, Invitrogen). After 3x 30 min washing steps (40mM Phosphate Buffer, 1mM EDTA, 5% SDS) the membrane was exposed to a phosphoscreen (BAS-IP SR2040, Fujifilm) and developed after 24 hours (FLA 5100R, Fujifilm). The Southern probe used for screening resistant clones after RMCE was kindly provided by Haydn Prosser. The probe mapped external to the targeting construct (*ROSA26* gene locus) was 400 bp in size. It was excised using KpnI/BglII, gel-purified, and 25 ng used for labeling and Southern blot screening.

### PCR-based screen

Human specific primer pairs used for validating the human BAC constructs and screening positive ES clones after RMCE were designed using Primer-BLAST (www.ncbi.nlm.nih.gov/tools/primer-blast/). The PCR conditions used were as follows: 1x DreamTaq Green Buffer (supplied as 10x stock by manufacturer), 0.2 mM dNTPs, 0.8 μg/μl BSA, 1 μM of each primer, 0.05 U/μl DreamTaq Polymerase, and either 150 ng BAC or 300 μg genomic template DNA. In most cases the genomic DNA had to be digested by EcoRI, EcoRV, KpnI or ScaI restriction enzymes before being used as a successful template for PCR reaction. PCR conditions were: 95° C for 3 min, 45 cycles at 95° C for 30 s, 60° C for 30 s, 72° C for 210 s, and 1 cycle of 72° C for 10 min. The primer sequence list is provided in **Supplementary Table 1**.

### Immunofluorescence

Mice were anesthetized intraperitoneally with Avertin (Sigma Aldrich) and perfused transcardially with 4% paraformaldehyde. Brains were post-fixed overnight at 4° C, cryoprotected in 30% sucrose, and frozen on dry ice. 40 µm sections were collected using a cryostat and stained with primary antibody overnight at 4° C (1:1000 rabbit α-GFP, Life Technologies) and incubated with secondary antibodies (2h at room temperature, goat IgG A647, Molecular Probes/Invitrogen) and DAPI. Confocal microscopy was performed with a TCS-SP5 Leica Laser Scanning System. The images were processed using ImageJ software (NIH).

### Protein extraction and luciferase assays

After perfusion with saline fresh brains were frozen on dry ice and stored at −80° C. For protein extraction, the whole brain of postnatal day 6 (P6) animals or dorsal raphe tissue punches (adult animals) were homogenized with Passive Lysis Buffer (Promega) and proteinase inhibitors (cOmplete ULTRA Tablets, Mini, EDTA-free, EASYpack, Roche) using an electrical homogenizer (POLYTRON PT 1200 E Manual Disperser, Kinematica). Tissue was further disrupted using a syringe and pipetting. The whole procedure was performed on ice. Samples were then shaken in a thermomixer at 1000 rpm and 4° C in a cold room for 1 hour, centrifuged, and supernatant collected. Samples were stored at −80° C until assaying. Luciferase assays (Firefly and Renilla) were done on the protein extracts according to manufacturer instructions (Dual-Luciferase Reporter Assay System, Promega). The signal was normalized to the amount of protein in the sample as determined by Pierce BCA Protein Assay Kit (Thermo Fisher Scientific) according to manufacturer instructions.

### Statistical analysis

In general, analysis of luciferase signal was performed using *Prism4* software (GraphPad). The effect of genotype and sex was assessed by two-way ANOVA. In case of significance ANOVA was followed by Tukey’s multiple comparison *post hoc* test to compare individual genotypes.

## Results

### 5-HTT-LPR-16A line creation and analysis

We took advantage of recombination-mediated cassette exchange (RMCE; Prosser et al., 2008) at the mouse *Rosa26* gene locus to create mouse lines harbouring a single copy insertion of the human 5-HTT gene locus with either the S or L promoter variant and carrying cistrons expressing both fluorescent and luciferase reporter proteins (Fig. 1A). We used recombineering to truncate a 5-HTT-containing human bacterial artificial chromosome (BAC) to include a 62 kb region containing the *5-HTT* gene, insert a neomycin selection cassette into the backbone, and add yellow fluorescent (YFP) and firefly luciferase (fLuc) protein open reading frames at the stop codon of the human 5-HTT via viral-derived 2A self-cleaving protein linkers (see Materials & Methods). The sequence of the inserted BAC was confirmed to be identical to the mouse reference sequence using Solexa deep sequencing (**Supplementary Fig. 1**). PCR of the 5-HTT-LPR region identified a human BAC contained the long, 16A polymorphism. Co-electroporation of the modified BAC with a Cre recombinase expressing plasmid (pCAGGS-Cre) allowed for the neomycin-resistance selection of RMCE-mediated insertion of the BAC into the mouse genome in an ES cell line engineered to carry LoxP and Lox511 sites at the *Rosa26* locus (Prosser et al., 2008). Selection and purification identified several ES cell clones carrying the modified human *5-HTT* locus correctly inserted at the mouse *Rosa26* locus as detected by Southern blot (Fig. 1B). RMCE is prone to the induction of inversions and/or deletions in the targeting construct (Prosser et al., 2008). Therefore, the integrity of the successfully targeted allele was confirmed by performing PCR assays on mouse genomic DNA from tail biopsy with a set of 44 human-specific primer pairs tiling the BAC at 100-500 bp intervals. Electrophoretic analysis of the resulting products allowed us to exclude several positive clones carrying large deletions (data not shown, Fig. 1C). Confirmed ES clones were then injected into mouse blastocysts to obtain chimeric offspring with at least one of them demonstrating germline transmission of the human 5-HTT-LPR-16A transgene. Analysis of brains of heterozygous transgene positive offspring revealed expression of YFP in sections from the dorsal raphe (Fig. 1D) as well as expression of firefly luciferase activity in tissue punches from the same region (Fig. 1E; one-tailed t-test, t_(3)_ = 2.758, P = 0.035 as compared to wild-type animals).

**Figure 1.**
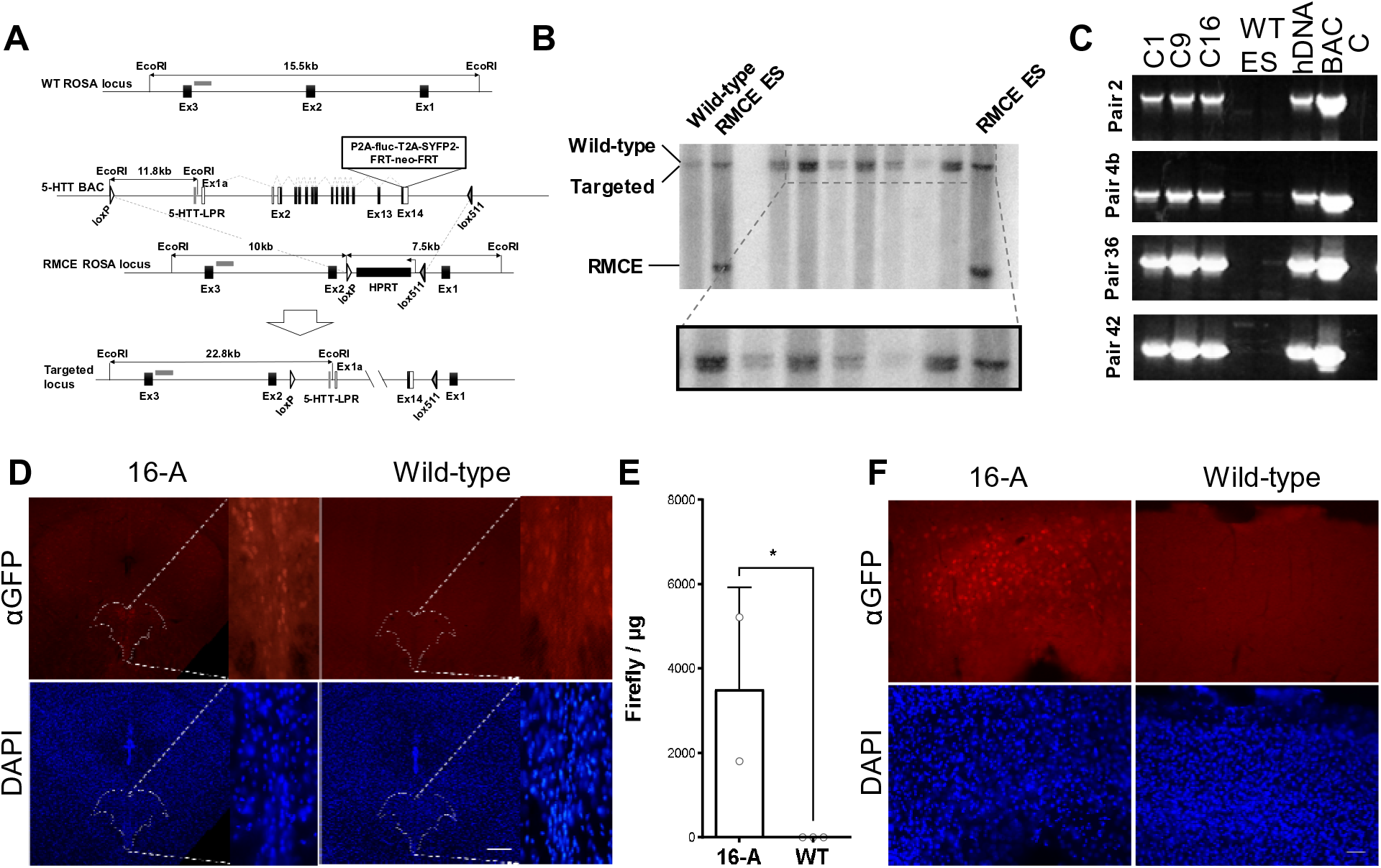
Generation of a mouse line carrying a single-copy human 5-HTT transgene. **(A-C)** The targeting vector was obtained by inserting a P2A-hLuc-T2A-SYFP2-FRT-Neo-FRT cassette at the stop codon of the human *5-HTT* gene within a 62 kb human genomic fragment (BAC). Cre/LoxP recombination-mediated cassette exchange (RMCE) was then used to insert a single copy of the targeting vector into an acceptor site engineered at the *Rosa26* in mouse ES cells. Proper ES cell targeting was verified by Southern Blot (**B**) transgene integrity was confirmed using a set of overlapping PCR probes spanning the region (**C**). (**D**) Expression of the human 5-HTT in the mouse brain as detected by the distribution of SYFP immunofluorescence in coronal brain sections from 5-HTT-LPR-16A mice (n=2). Positive cell bodies are observed in the dorsal raphe nucleus (dashed line) and at lower levels outside the raphe in transgenic mice (intensity in the raphe is 40% higher). Background staining is seen in littermate wild-type (WT) controls. Scale bar: 200 µm. (**E**) Luciferase assays on adult whole brain protein extracts confirmed transgene expression in 16A (n=2), but not WT controls (n=3; data shown as mean±SD). (**F**) Ectopic expression of the human 5-HTT in the mouse brain as detected by the distribution of SYFP immunofluorescence in coronal brain sections from 5-HTT-LPR-16A observed in the cortex (scale bar: 50 µm).

### 5-HTT-LPR-14A line creation and analysis

To determine the effect of the 5-HTT-LPR polymorphism on the expression of 5-HTT we constructed a mouse line in which the 16A sequence was mutated to 14A. Unfortunately, several bacterial recombineering strategies were unsuccessful at replacing the 16A variant by the 14A variant without modifying the surrounding genomic DNA (see **Materials & Methods**). Instead, we settled on a two-step, positive/negative-selection recombineering technique (Muyrers et al., 2000) in which we first replaced the 5-HTT-LPR in our 16A modified human BAC with an AMP-SacB negative selection cassette. Next, we introduced a Zeomycin (Zeo) positive selection gene flanked by rox site specific recombinase recognition sites (Anastassiadis et al., 2009) upstream of the 14A variant within a smaller targeting plasmid and used recombineering to replace AMP-SacB within the larger trimmed human BAC with the 5-HTT-LPR-14A sequence. Finally, the Zeo selection gene was removed by Dre-mediated excision leaving behind a 32 bp rox site 116 bp upstream of 5-HTT-LPR (Fig. 2A). This strategy resulted in the creation of two BAC constructs identical except for the 5-HTT-LPR 16A/14A site, but that differed from the endogenous human locus by the presence of the rox site. Both constructs were used for RMCE targeting in ES cells, resulting in several positive clones as assessed by Southern blotting (Fig. 2B). The integrity of the BACs within the resulting ES cell lines was confirmed by the previously described nested PCR-method (Fig. 2C). Confirmed clones were then used to generate and establish the 16A-Rox and 14A-Rox mouse lines.

**Figure 2.**
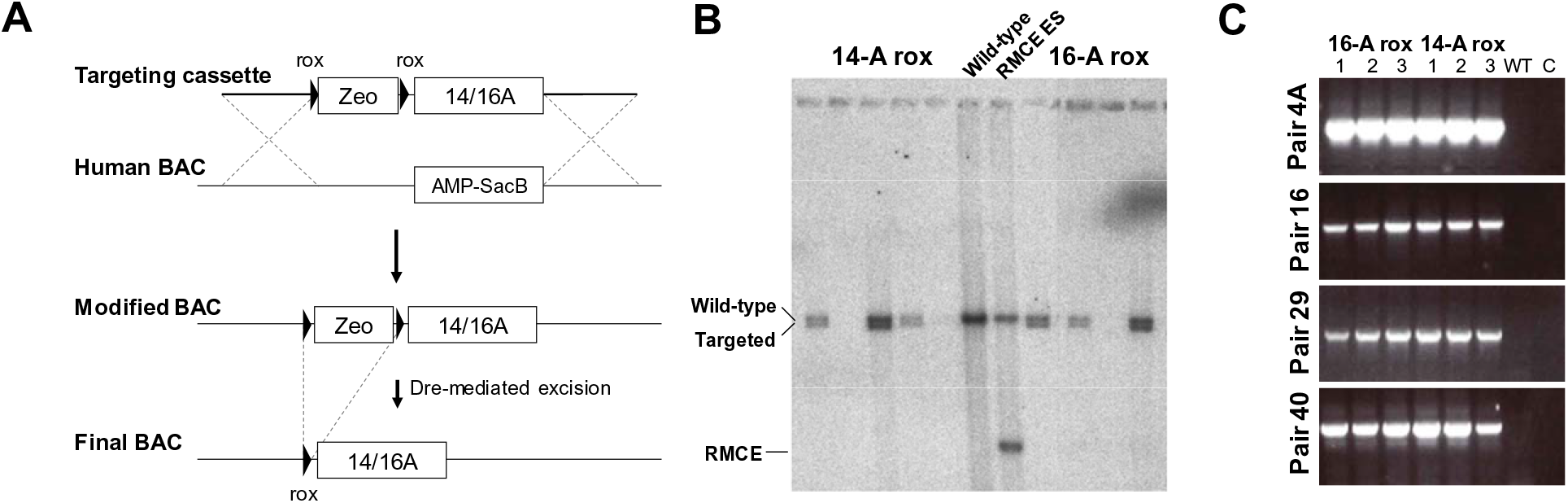
Generation of 5-HTT-LPR-16A-Rox and 14A-Rox mouse lines. **(A)** Generation of mice bearing the human 5-HTT-LPR Short (14A) allele. Cassettes carrying either the 14A or 16A polymorphisms with a rox-flanked Zeo-selection cassette inserted 116 bp upstream were used to replace the AMP-SacB cassette inserted into a 5-HTT- LPR containing BAC. Following selection of the correctly inserted polymorphic cassettes the Zeo selection gene was removed by Dre-mediated recombination. This procedure left a 32 bp rox site upstream of 5-HTT-LPR. The 16A-Rox and 14A-Rox constructs were separately targeted to a mouse *Rosa26* acceptor locus using Cre-dependent RMCE to produce single-copy insertions of the human 5-HTT locus at the same mouse genome location. (**B-C**) Targeting of the *Rosa26* acceptor locus via RMCE and Southern blot screening using a 3’ radiolabeled probe. Positive ES clones were picked for expansion and further screening with the same PCR-based approach used to test transgene integrity above. Representative samples of 4 primer pairs are shown.

To determine whether the 16A and 14A variants were associated with differential expression of 5-HTT in the context of mouse brain development, we quantified firefly luciferase activity in brain samples taken from either postnatal day 6 (P6) pups or adults of both lines. Two-way ANOVA analysis revealed no main effect of genotype (F_1,25_ = 1.89, P = 0.181), but a significant main effect of sex (F_1,25_ = 4.729, P = 0.039), and no interaction between the two (F_1,25_ = 0.392, P = 0.537), suggesting that the 5-HTT-LPR polymorphism does not affect 5-HTT expression during development, at least as assessed by firefly luciferase activity at this developmental stage (Fig. 3A). In adult animals we were able to use tissue punches to reliably excise tissue from the dorsal raphe region of the brainstem where serotonin neurons have their cell bodies. To control for possible variation in the recovery of serotonin cell bodies within the excised tissue we crossed the 16A-Rox and 14A-Rox lines to a *Tph2*::rLuc-SCFP transgenic mouse line in which Renilla luciferase is expressed selectively in serotonin neurons (Mlinar et al., 2016) and normalized our firefly luciferase measurements to Renilla luciferase within each sample. As expected, firefly luciferase activity was detected only in brain samples from mice carrying either 16A-Rox or 14A-Rox, but not in those carrying only *Tph2*::rLuc-SCFP (**Supplementary Fig. 2A**, one-way ANOVA. F_3,7_ = 36.36, P = 0.0001 followed by Tukey post-hoc test). Conversely, Renilla luciferase activity was detected only in brain samples from mice carrying the *Tph2*::rLuc-SCFP transgene (**Supplementary Fig. 2B**, one-way ANOVA. F_3,7_ = 44.01, P < 0.0001, followed by Tukey post-hoc test). Two-way ANOVA analysis of firefly luciferase activity in dorsal raphe tissue punches from adult double transgenic mice (Fig. 3B) showed no main effect of genotype (F_1,8_ = 0.8805, P = 0.3755), sex (F_1,8_ = 0.9958, P = 0.3476), nor interaction between the two (F_1,8_ = 0.4039, P = 0.5428). Similarly, firefly luciferase activity normalized to Renilla luciferase signal (Fig. 3C) in another cohort of animals showed no main effect of genotype (F_1,26_ = 2.482, P = 0.127), sex (F_1,26_ = 1.613, P = 0.215), nor interaction between the two (F_1,26_ = 2.342, P = 0.138). These findings suggest that the 5-HTT-LPR polymorphism does not affect 5-HTT gene expression in either early postnatal development or adulthood in the mouse.

**Figure 3.**
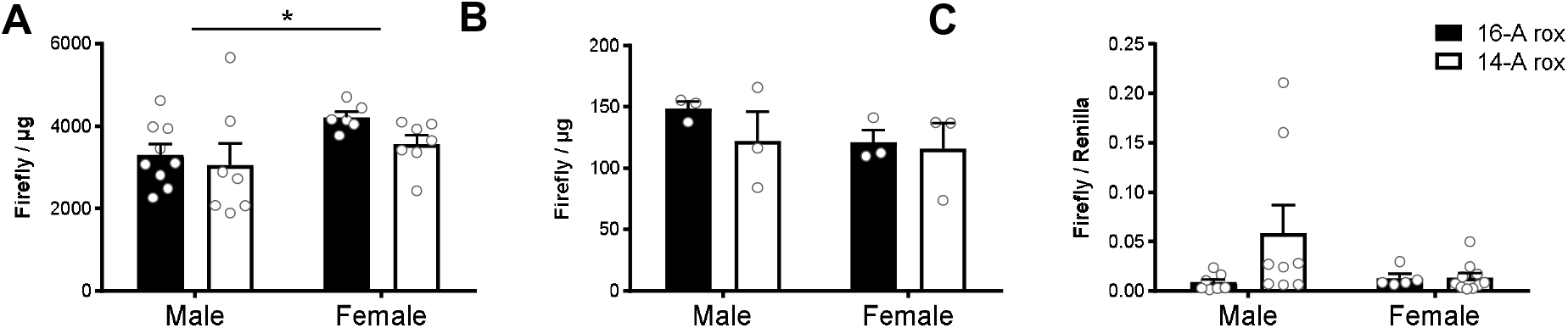
Transgene expression in 16A-Rox and 14A-Rox lines. **(A)** Firefly luciferase activity in whole brain samples from 16A-Rox (n = 15, 9 males, 6 females) and 14A-Rox (n = 14, 7 ales, 7 females) at postnatal day 6. No significant effect of genotype was detected, but female mice showed significantly greater transgene expression than males. (**B**) Luciferase activity in tissue punches from the dorsal raphe of adult (postnatal day 60) 16A-Rox;*Tph2*::rLuc-SCFP and 14A-Rox;*Tph2*::rLuc-SCFP mice revealed no significant genotype or sex effects (16A-Rox: n = 6, 3 males, 3 females; 14A-Rox: n = 6, 3 males, 3 females). (**C**) Relative luciferase activity was used to control for potential inconsistencies in the recovery of serotonin cell bodies in the raphe tissue punches (16A-Rox: n = 12, 7 males, 5 females; and 14A-Rox: n = 19, 8 males, 11 females; data shown as mean ± SEM).

## Discussion

Our findings are consistent with data suggesting that the 5-HTT-LPR polymorphism does not affect 5-HTT expression in adult human brain (Parsey et al., 2006; Shioe et al., 2003). However, several technical issues concerning our ability to faithfully model human gene expression in the mouse call into question this conclusion. First, although we were able to detect expression of the human 5-HTT gene from our BAC transgenic locus in the 16A line (Fig. 1E), we also detected low levels of ubiquitous transgene expression (Fig. 1F) suggesting that either our transgenic construct is missing important human regulatory sequences or that the mouse host is missing critical transcription factors required to suppress ectopic expression of the human locus. It is possible that these missing regulatory mechanisms are necessary for the polymorphism to impact gene expression. Second, our data suggest that the Rox site left upstream of the 5-HTT-LPR as a result of our recombineering strategy significantly interfered with expression of the human transgene. We were unable to detect SYFP in tissue sections from either 16A-Rox or 14A-Rox mice and absolute levels of luciferase activity from the Rox-containing lines was several orders of magnitude less that that detected in similar samples from the 16A line (see Fig. 1E and Fig. 3B). Thus, it remains possible that differences in expression between the 16A-Rox and 14A-Rox lines could not be detected under conditions in which critical regulatory mechanisms were missing and exogenous sequences interfered with robust expression of the human transgene.

Our study serves as a cautionary note for future studies aimed at studying human transgene regulation in the context of the living mouse. Despite taking extensive precautions to ensure that we were comparing the effect of the human variant within the context of an otherwise identical single-copy transgene engineered at the same genomic location, we encountered several problems that led to both ectopic and hypomorphic expression of the human transgene in the mouse brain. As advances in human genetics rapidly uncovers putative genetic variants implicated in disease susceptibility and researchers are increasingly seeking to use the mouse as a platform to uncover their molecular and physiological impact, our study suggests that care must be taken to ensure that the human variants are studied within a sufficiently faithful context.

## Acknowledgments

We thank Olga Ermakova for establishing the RMCE technology in the laboratory and for help with Southern blot analysis, Haydn Prosser for generously sharing reagents prior to publication, Vladimir Benes (EMBL Genomic Core Facility) for advice and for sequencing the 16A human BAC. This work was supported by funding from EMBL.

