## Supplementary Figure 1 for "Mouse model of the human serotonin transporter-linked polymorphic region"

### Supplementary Figures and Tables

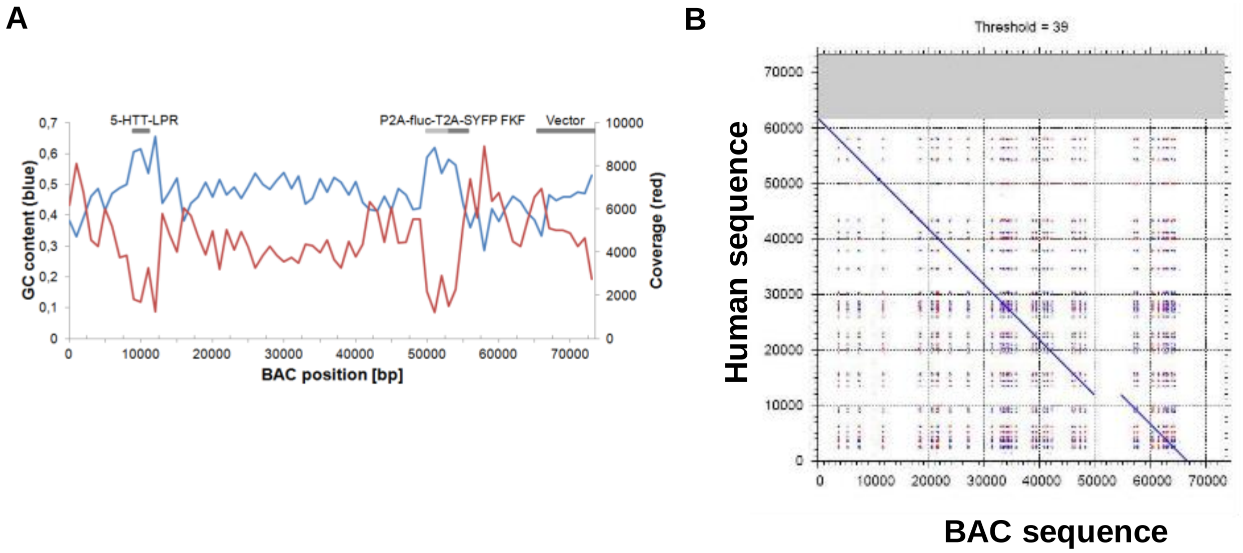


***Supplementary Figure 1.*** **Sequence of the modified 16A BAC.** (**A**) GC content (blue) and coverage (red) of the human 5-HTT-LPR-16A targeting construct modified to co-express firefly luciferase (fLuc) and SYFP. (**B**) Comparison of the sequence with the human reference sequence showing high sequence identity (diagonal line). The gap in the diagonal line results from the co-expression cassettes at the 3' end of the 5-HTT open reading frame in the targeting construct.

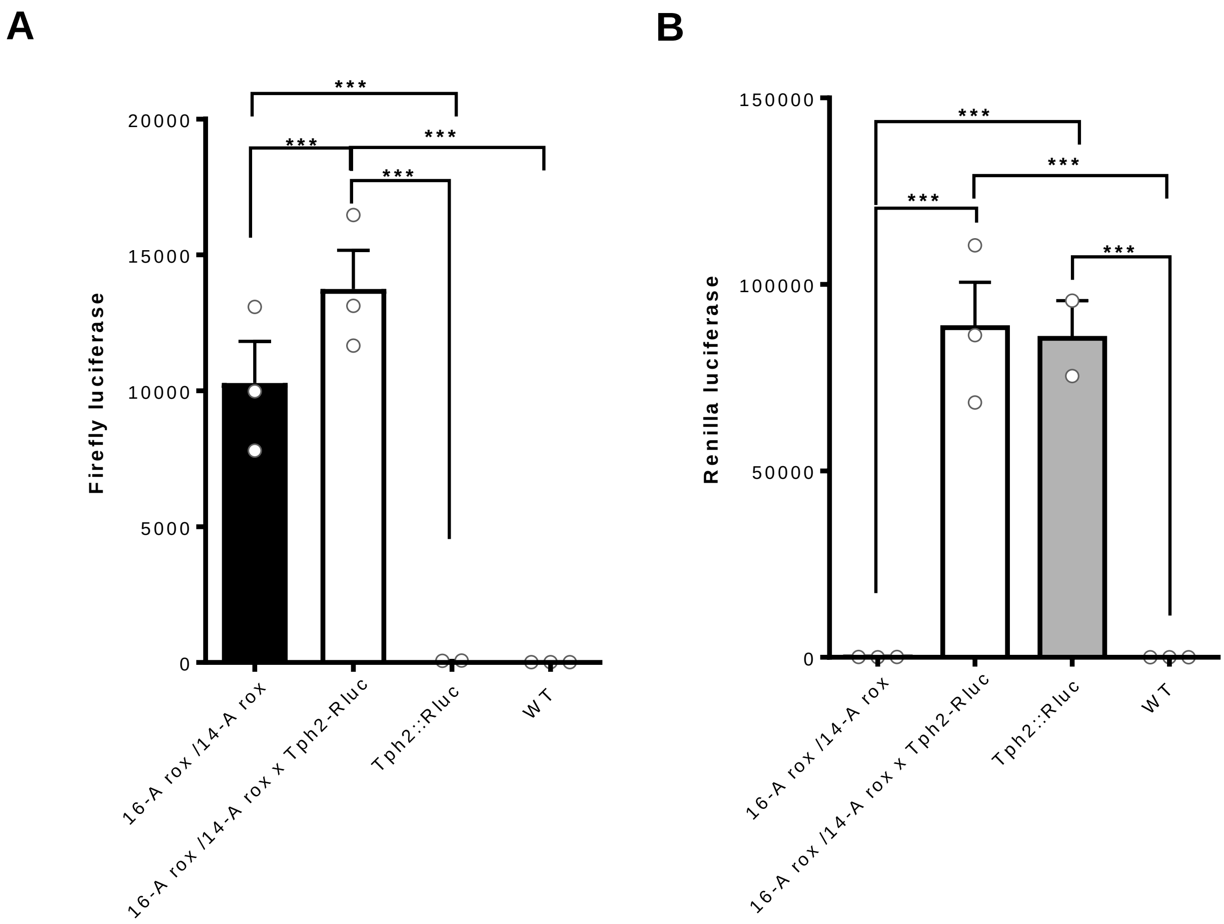


***Supplementary Figure 2.*** **Independent detection of Firefly and Renilla luciferase signal in the same sample.** Absence of cross-contamination between Firefly and Renilla in the dual luciferase assay system. (**A**) Firefly luciferase signal was observed in all 16A-Rox and 14A-Rox containing mouse lines, but not in those carrying only *Tph2*::rLuc-SCFP. (**B**) Renilla luciferase signal was observed in all *Tph2*::rLuc-SCFP containing mouse lines, but not in those carrying only 16A-Rox or 14A-Rox (***p < 0.005, one-way ANOVA, Bonferroni post-hoc, compared to WT group; data shown as mean±SEM).

***Supplementary Table 1.*** Human specific PCR primer sets used to confirm transgene integrity.

| ***Pair no.*** | **Size of product (kb)** | **Primer name** | **Oligo Sequence 5'-> 3'** |
|---|---|---|---|
| *1* | 2.4 | WG hBAC 1F | AAAAGGGCTTCCCTCTTCAG |
| | | WG hBAC 1R | GTTCTAGCAGCCAAGGATGG |
| *2* | 2.5 | WG hBAC 2F | TCCACAAGCTCGGGAAGACCAGG |
| | | WG hBAC 2R | CCCCTGGGAGCCAACGGAGTA |
| *3* | 2.4 | WG hBAC 3F | TGCACACATTGACAGGAACA |
| | | WG hBAC 3R | GGCAGTTTTCTATGGGCTGA |
| *4a* | 1.0 | WG hBAC 4Fa | TGCGTCTGGTGTAACATGGAGCTGA |
| | | WG hBAC 4Fb | GAAGGGAATCCCCTAGGGAGGAAGT |
| *4b* | 1.5 | WG hBAC 4Ra | GTGGCGCCAAGGTCGTCTGGC |
| | | WG hBAC 4Rb | TGGGGGAGGAACAGATCAGGTTCAG |
| *5* | 2.5 | WG hBAC 5F | TGTGGTGAGGCTGCTGCCGTA |
| | | WG hBAC 5R | AGGAGGGGTACAGGGAGTTGCT |
| *6* | 1.9 | WG hBAC 6F | AGCCGTCCTCCGCTTTGGC |
| | | WG hBAC 6R | TGCGTCACTTTGAGGCGAATAAACT |
| *L* | 0.8 | WG hBAC linkerR | AACGTGGGTTCGAGGCGGAGA |
| | | WG hBAC linkerF | GCACTGTGCTCCTTTTTGACGCA |
| *8* | 1.6 | WG hBAC 8F | GCAGGTGGGTCCGCTTTTCC |
| | | WG hBAC 8R | GCGGACATCCAGGGTGTTTGGG |
| *9* | 2.3 | WG hBAC 9F | CCCCTCAACACTCTGGTGATCCA |
| | | WG hBAC 9R | GTGGGTGAGCTAGAGCCAGCCA |
| *10* | 2.0 | WG hBAC 10F | TCTGGGCGAGCACTTCTCAGCA |
| | | WG hBAC 10R | CGAGGCTGGGAAAGAGACTTGAGA |
| *11* | 2.5 | WG hBAC 11F | AGCTGTGTGTGGACATGTTCCCATG |
| | | WG hBAC 11R | CCTGCCTGGGAGAGGATAGCCC |
| *12* | 2.5 | WG hBAC 12F | GTGGGCCTCAGTTTCCCTGCTAG |
| | | WG hBAC 12R | ATAGCCAGGAGCAACCCGTACCTG |
| *13* | 2.5 | WG hBAC 13F | TTACTGCTTCGGGCGGCACCA |
| | | WG hBAC 13R | TTTATGCTACTGCCTGGCCTTGGC |
| *14* | 1.8 | WG hBAC 14F | CCTTCCGTAGACCCTCTGGGCC |
| | | WG hBAC 14R | AACGGCACTGCTGCTCACCAT |
| *15* | 1.4 | WG hBAC 15F | TCCAGTGTCATCTCAGCTAGGCAG |
| | | WG hBAC 15R | CTCCCGCACCAGGACTTGGA |
| *16* | 1.4 | WG hBAC 16F | TCCAGCCTGTGCAAACCTTGGTGAT |
| | | WG hBAC 16R | GCTTCACAACCCGGAGAGCCTC |
| *17* | 2.3 | WG hBAC 17F | ACACGGCACTCTATCCCAGCG |
| | | WG hBAC 17R | CAGCTGCAACTCTCTGTGAGTCACC |
| *18* | 2.4 | WG hBAC 18F | TTGGAGGAAGGCCATCACGAGAACA |
| | | WG hBAC 18R | AAGCACATTTGGCCAACACCCTGG |
| *19* | 2.3 | WG hBAC 19F | TACCTCAAGTGCTCCACGGCCT |
| | | WG hBAC 19R | GGACCCCAAAGCCCGGACCAA |

| *Pair no.* | **Size of product (kb)** | **Primer name** | **Oligo Sequence 5'-> 3'** |
|---|---|---|---|
| *20* | 1.8 | WG hBAC 20F | AGGAGTGGCGACCCTGTTGGT |
| | | WG hBAC 20R | CACTGCAGCCTTAACCTCCCGG |
| *21* | 1.1 | WG hBAC 21F | CAGGAGGATTGCTTGAGCCCGG |
| | | WG hBAC 21R | GCGTCTTTGGCCACCTCAGAC |
| *22* | 2.1 | WG hBAC 22F | GCGGCCCCTTGGGTTTTCCC |
| | | WG hBAC 22R | CAGTTCCCCGGCTCGCTGG |
| *23* | 0.8 | WG hBAC 23F | AATTATGGCGCCTACAGGCCGG |
| | | WG hBAC 23R | GCGGTAAAATGCTGACAGCCCCTG |
| *24* | 0.8 | WG hbAC 24F | GGCTTGGAGGGGGTGATCACG |
| | | WG hBAC 24R | TTGAACACCTGCTGTGGACACGA |
| *25* | 1.7 | WG hBAC 25F | GGCTGTCAGCATTTTACCGCAGA |
| | | WG hBAC 25R | AGCCCTGTGTGCAACCCAAAATGT |
| *26* | 2.4 | WG hBAC 26F | ACCTCCCGATGATGGGTCTGTAAAA |
| | | WG hBAC 26R | AGGGCCCCCATGAAGAGCATAGC |
| *27* | 1.6 | WG hBAC 27F | AGTGGTTATAGTTCTGGCACTCCCA |
| | | WG hBAC 27R | TCTGGTAGGATGGGCTAGGCGA |
| *28* | 2.3 | WG hBAC 28F | TCTTGTCTCTGTGACACTGCATGTG |
| | | WG hBAC 28R | TTACCAACTTCTGTACCCAAGCTGC |
| *29* | 1.8 | WG hBAC 29F | ATGATGAGGACCTGTGGCAGGC |
| | | WG hBAC 29R | AATTAGCCCACCTCCGTTCTCAGGC |
| *30* | 2.0 | WG hBAC 30F | GTACCTTGGGAGGTAAATGGGCAGG |
| | | WG hBAC 30R | CGCAAGACTTCCAGACCGTGGGT |
| *31* | 2.1 | WG hBAC 31F | ACGCCGAGGTGGATGGGTCA |
| | | WG hBAC 31R | TGCGGCCACAAAGGGCTTGG |
| *32* | 2.4 | WG hBAC 32F | TTTCAAAGCCTCTGCAGTGTGGCT |
| | | WG hBAC 32R | TTGCCGTGTTCCAGCACGACG |
| *33* | 2.4 | WG hBAC 33F | GCGCCCCACTGTCTAAGGAGGT |
| | | WG hBAC 33R | TTACGGTATCGCCGCTCCCGA |
| *34* | 2.1 | WG hBAC 34F | AGAAACGCGGGCGTATTGGCC |
| | | WG hBAC 34R | ATCTTCCTTTCCTGATGCCACAGCA |
| *35* | 1.8 | WG hBAC 35F | CCAGTTCTGATGAGGCACGCC |
| | | WG hBAC 35R | GGGTGGGGAGAATCCATATCCGA |
| *36* | 1.3 | WG hBAC 36F | TCCAGACAGCAGAGGAAACATCCT |
| | | WG hBAC 36R | ATTAGCTTGGTCCCCAGGTTTGTCT |
| *37* | 1.0 | WG hBAC 37F | AGTGAGCCAAACGTTCCACTGCA |
| | | WG hBAC 37R | TGATGCCATCTATTCTCTCCCCCAA |
| *38* | 1.6 | WG hBAC 38F | TGCCAAGCTGCTCTACCACCAATG |
| | | WG hBAC 38R | GGCAGGAGGTCTGTTAGGAACTGGA |
| *39* | 1.9 | WG hBAC 39F | CTTTTTGGCACCGGGACCGGT |
| | | WG hBAC 39R | GGTCGAGGCTGTAGTGAGCTGTGA |
| *40* | 1.1 | WG hBAC 40F | ACTGCTGCATGACTGTCAACTCATC |
| | | WG hBAC 40R | ATCCAGTGAGAGGGCTGAGGATCA |

| *Pair no.* | Size of product (kb) | Primer name | Oligo Sequence 5'-> 3' |
|---|---|---|---|
| *41* | 1.4 | WG hBAC 41F | TTGCCCAGGCGGTTCTCCTG |
| | | WG hBAC 41R | TGTCGCCCAGGCTAGAGTGC |
| *42* | 1.2 | WG hBAC 42F | ATTCCTGGTGCATCTGTGGAACACG |
| | | WG hBAC 42R | GGAACTTCTCCCTCCCCTCTATGGC |
| *43* | 1.3 | WG hBAC 43F | AATACCGTTGCTTCGCATCCTCATC |
| | | WG hBAC 43R | TTCCAGTTGCCCCACATCTTGCT |
| *44* | 1.2 | WG hBAC 44F | AGGCAGGCGAAATTGCTTGAGCC |
| | | WG hBAC 44R | TCCACGGGTTCAAATGGGGCAG |
